# Interleukin-1 receptor antagonist mediates type I interferon-driven susceptibility to *Mycobacterium tuberculosis*

**DOI:** 10.1101/389288

**Authors:** Daisy X. Ji, Katherine J. Chen, Naofumi Mukaida, Igor Kramnik, K. Heran Darwin, Russell E. Vance

**Affiliations:** Division of Immunology and Pathogenesis, Department of Molecular and Cell Biology, University of California, Berkeley, CA 94720 USA; Division of Molecular Bioregulation, Cancer Research Institute, Kanazawa University, Kakuma-machi, Kanazawa 920-1192, Japan; The National Emerging Infectious Diseases Laboratory, Department of Medicine (Pulmonary Center), and Department of Microbiology, Boston University School of Medicine, Boston, MA 02118 USA; Department of Microbiology, New York University School of Medicine, New York, New York, 10016 USA; Cancer Research Laboratory, University of California, Berkeley, CA 94720 USA; Howard Hughes Medical Institute, University of California, Berkeley, CA 94720 USA

## Abstract

The bacterium *Mycobacterium tuberculosis* (*Mtb*) causes tuberculosis (TB) and is responsible for more human mortality than any other single pathogen^1^. Although ~1.7 billion people are infected with *Mtb*^2^, most infections are asymptomatic. Progression to active disease occurs in ~10% of infected individuals and is predicted by an elevated type I interferon (IFN) response^3–8^. Type I IFNs are vital for antiviral immunity, but whether or how they mediate susceptibility to *Mtb* has been difficult to study, in part because the standard C57BL/6 (B6) mouse model does not recapitulate the IFN-driven disease that appears to occur in humans^3–5,8^. Here we examined B6. *Sst1^S^* congenic mice that carry the C3H “sensitive” allele of the *Sst1* locus that renders them highly susceptible to *Mtb* infections^9,10^. We found that B6.Sst1^S^ mice exhibit markedly increased type I IFN signaling, and that type I IFNs were required for the enhanced susceptibility of B6. *Sst1^S^* mice to *Mtb*. Type I IFNs affect the expression of hundreds of genes, several of which have previously been implicated in susceptibility to bacterial infections^11,12^. Nevertheless, we found that heterozygous deficiency in just a single IFN target gene, IL-1 receptor antagonist (IL-1Ra), is sufficient to reverse IFN-driven susceptibility to *Mtb*. As even a partial reduction in IL-1Ra levels led to significant protection, we hypothesized that IL-1Ra may be a plausible target for host-directed anti-TB therapy. Indeed, antibody-mediated neutralization of IL-1Ra provided therapeutic benefit to Mtb-infected B6. *Sst1^S^* mice. Our results illustrate how the diversity of inbred mouse strains can be exploited to better model human TB, and demonstrate that IL-1Ra is an important mediator of type I IFN-driven susceptibility to *Mtb* infections *in vivo*.

## Introduction

*Mycobacterium tuberculosis* (*Mtb*) infects approximately one quarter of all humans worldwide^2^, with highly diverse outcomes ranging from asymptomatic lung granulomas to lethal disseminated disease. Active TB disease is characterized by the uncontrolled replication of bacteria and pathological inflammation in the lungs and other organs, and arises in approximately 10% of infected HIV-negative individuals. There is no vaccine that reliably protects against pulmonary TB, and although antibiotics can be curative, the long (≥6-month) course of treatment and increasing prevalence of multi-drug resistant *Mtb* infections has spurred a search for alternative therapeutic approaches^13,14^. Since only individuals with active TB readily transmit infection, and as humans are the only natural reservoir of *Mtb*, a favored strategy to contain the TB epidemic is to identify and treat latently infected individuals likely to progress to active disease^1,3^. Identification of such individuals is challenging, but recent studies have demonstrated that an enhanced type I interferon (IFN) signature correlates with active TB^5,6,8^ and can predict progression to active TB up to 18 months prior to diagnosis^3,4^. A partial loss-of-function polymorphism in the type I IFN receptor (IFNAR1) is associated with resistance to TB in humans, suggesting that elevated levels of type I IFNs not only predict but may even be causally linked to TB progression^15^. In addition, numerous animal studies have demonstrated causal roles for type I IFNs in susceptibility to *Mtb*^6,7,16,17^ and other bacterial infections^11,12^.

Given strong evidence that type I IFNs confer susceptibility to human TB, two major remaining challenges are: (1) to determine the mechanisms by which type I IFNs mediate susceptibility to active TB, and (2) to exploit this knowledge to develop interventions that can reverse the susceptibility. Mechanistic studies and initial trials of possible therapeutic interventions require a robust animal model. However, the most commonly used animal model, the C57BL/6 (B6) mouse, does not robustly recapitulate the interferon-driven TB susceptibility seen in humans. B6 *Ifnar^−/−^* mice show mild resistance to *Mtb* in the spleen and variable but modest effects in the lungs^18−20^. To better model IFN-driven susceptibility, some investigators treat Mtb-infected B6 mice with poly-IC, a potent inducer of type I IFNs^21^. Such treatment dramatically increases susceptibility to *Mtb* in an *Ifnar-dependent* manner^21,22^, but because the IFN is induced artificially and not by *Mtb* itself, it is unclear if poly-IC mimics the course of IFN-driven disease in humans.

As an alternative approach, we sought to exploit the natural diversity of available inbred mouse strains. The 129 mouse strain shows clear IFN-driven susceptibility to *Mtb*^23^, but there are limited tools on this genetic background. We therefore turned to a previously described congenic mouse strain, B6.*Sst1^S^*, that carries the 10.7Mb ‘super susceptibility to tuberculosis 1’ region of mouse chromosome 1 from C3H on an otherwise B6 genetic background^9,10^. The B6. *Sst1^S^* mice exhibit marked susceptibility to aerosol TB infection^9,10^, however the mechanism by which the *Sst1^S^* locus confers susceptibility remains incompletely understood. Recent work has established that macrophages from B6.Sst1^S^ mice exhibit an enhanced type I IFN response^24,25^. Here we show that B6.*Sst1^S^* mice exhibit an exacerbated type I IFN response *in vivo* that causes susceptibility to *Mtb* infection. We exploit this model of type I IFN-driven TB disease to identify the interleukin-1 receptor antagonist (IL-1Ra) as a major driver of type I IFN-induced susceptibility to *Mtb*. Genetic or antibody-mediated reduction in IL-1Ra levels largely eliminates the IFN-induced susceptibility of B6.*Sst1^S^* mice. Our results suggest new therapeutic strategies for tuberculosis.

## Results

In agreement with previous work^24,25^, we found that bone marrow-derived macrophages (BMMs) from B6.*Sst1^S^* mice expressed higher levels of interferon-beta (*Ifnb*) and interferon-stimulated genes (ISGs) upon stimulation with TNF (Fig. S1). To determine if B6.*Sst1^S^* mice also exhibit an enhanced type I IFN signature *in vivo*, we measured *Ifnb* transcripts in the lungs of *M. tuberculosis-infected* mice. Indeed, B6.*Sst1^S^* mice exhibited higher levels of *Ifnb* transcripts as compared to B6 mice (Fig. 1a). To investigate whether this enhanced type I IFN signaling causes the susceptibility of B6.*Sst1^S^* mice to *Mtb*, we infected mice with *Mtb* and then treated with an IFNAR1-blocking antibody^26^ to inhibit type I IFN signaling. B6.*Sst1^S^* mice treated with the IFNAR1-blocking antibody showed significantly decreased bacterial burdens compared to those that only received an isotype control antibody (Fig. 1b). To provide genetic confirmation of this result, we crossed B6.*Sst1^S^* mice to B6.*Ifnar^−/−^* mice. *Ifnar* deficiency largely reversed the enhanced susceptibility of B6.*Sst1^S^* mice to *Mtb* infection. At 25 days post-infection, the bacterial burdens in the lungs of the B6. *Sst1^S^Ifnar^−/−^* mice were significantly lower than in the lungs of B6.*Sst1^S^* mice, and were similar to B6 mice (Fig. 1c). Infected B6.*Sst1^S^Ifnar^−/−^* mice also survived significantly longer than B6.*Sst1^S^* mice (Fig. 1d), though there are also clearly *Ifnar-independent* effects of the *Sst1^S^* locus that act at later time points. By contrast, and consistent with prior reports^18–20^, *Ifnar* deficiency had little or no effect on *Mtb* disease in wild-type B6 (*Sst1^R^*) mice. Recent data have suggested that the host protein STING is required for interferon induction to *Mtb*^27–31^. However, crossing B6.*Sst1^S^* mice to STING-deficient *Sting^gt/gt^* mice did not significantly reduce bacterial burdens at day 25 compared to B6.*Sst1^S^* mice (Fig. S2a). However, the B6. *Sst1^S^Sting^gt/gt^* mice did show a slight improvement in survival not seen in the B6 genetic background^30^ (Fig. S2b). Overall our data demonstrate that the *Sst1^S^* locus acts to increase type I IFN signaling *in vivo* and thereby exacerbate *Mtb* infection, particularly during the early phases of infection.

**Fig. 1.**
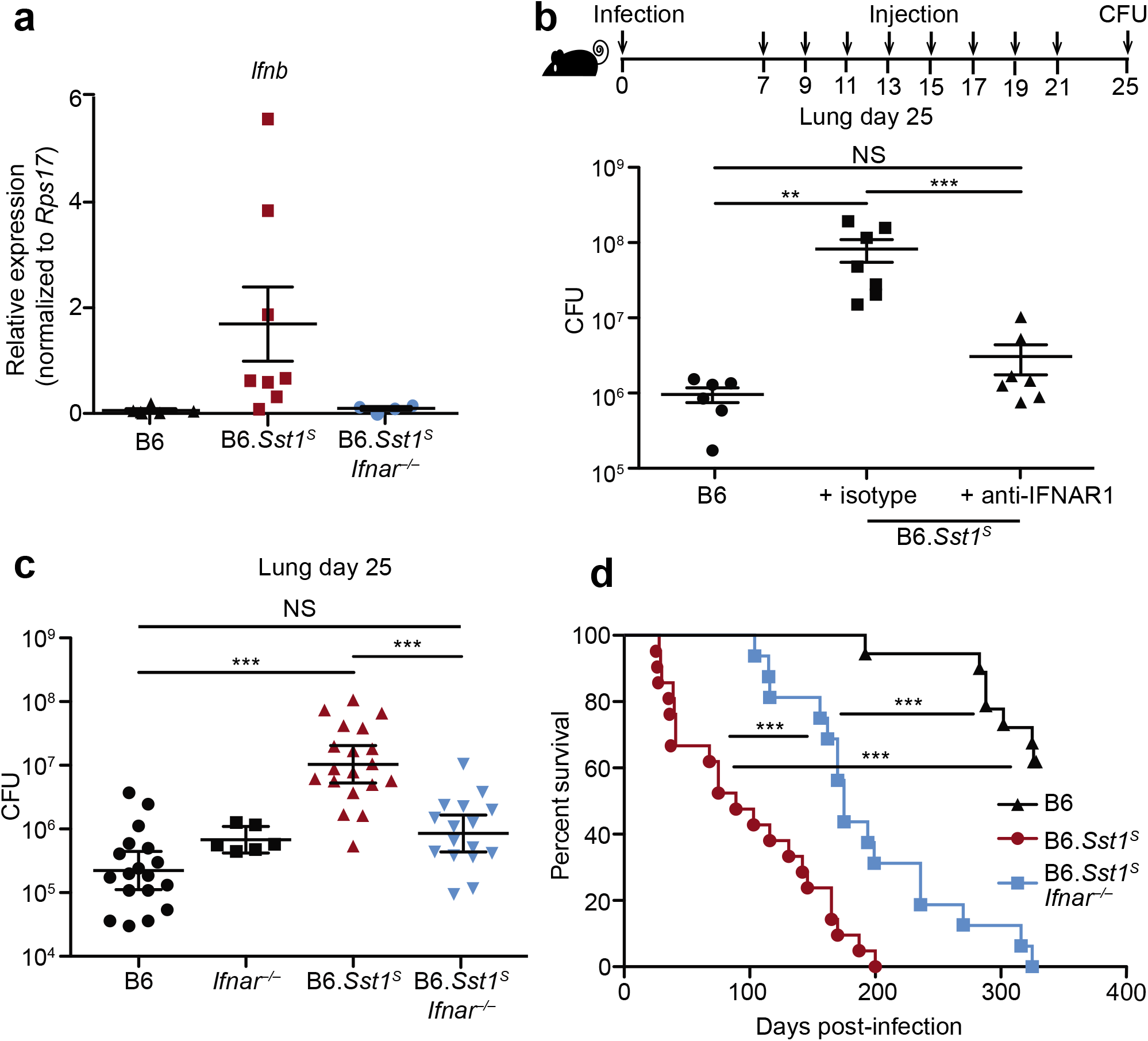
Type I IFN drives enhanced susceptibility of B6.*Sst1^S^* mice. **a**, Expression of *Ifnb* measured by qRTPCR relative to *Rps17* in Mtb-infected lungs at day 25. **b**, Lung bacterial burdens at day 25 from Mtb-infected mice treated with anti-IFNAR1 or isotype control antibody. **c**, Lung bacterial burdens at day 25, or **d**, survival, of Mtb-infected mice. For **a-c**, genotypes indicated on the x-axis. **c-d**, Combined results from three independent infections. All except B6 mice were bred in-house (**b-d**). Error bars are SEM. Analyzed with two-ended Mann-Whitney test (**b, c**) or Log-rank (Mantel-Cox) Test (**c**). Asterisk, *p* ≤ 0.05; two asterisks, *p* ≤ 0.01; three asterisks, *p* ≤ 0.001.

Type I IFN negatively regulates anti-bacterial immune responses via multiple mechanisms^6,7^, including through increased IL-10 levels^32^, decreased IFNγ signaling^33^, induction of cholesterol 25-hydroxylase (*Ch25h*)^34^, and/or decreased IL-1 levels^35,36^. We did not observe significant differences in IL-10 or IFNγ levels in the lung during *in vivo Mtb* infection (Fig. S3a,b). In addition, crossing B6.*Sst1^S^* mice to B6.*Ch25h^−/−^* mice did not alter day 25 lung bacterial burdens (Fig. S3c). Moreover, despite clear evidence that type I IFN and IL-1 counter-regulate each other^37^, the *Sst1^S^* locus did not appear to act to decrease the levels of IL-1 *in vivo*; in fact, we unexpectedly observed higher levels of both IL-1α and IL-1β in the lungs of B6.*Sst1^S^* mice at 25 days post-infection as compared to B6 mice (Fig. 2a,b). Other inflammatory cytokines, including TNF and CXCL1 were similarly elevated in the B6.*Sst1^S^* mice (Fig. S3d,e) as was the frequency of CD11b^+^Ly6G^+^ cells (neutrophils) in the lungs (Fig. S3f). The elevated inflammation in B6.*Sst1^S^* mice was a consequence of elevated type I IFNs, as inflammatory cytokines and neutrophils were reduced in B6.*Sst1^S^Ifnar^−/−^* mice, but the underlying mechanism was not apparent.

**Fig. 2.**
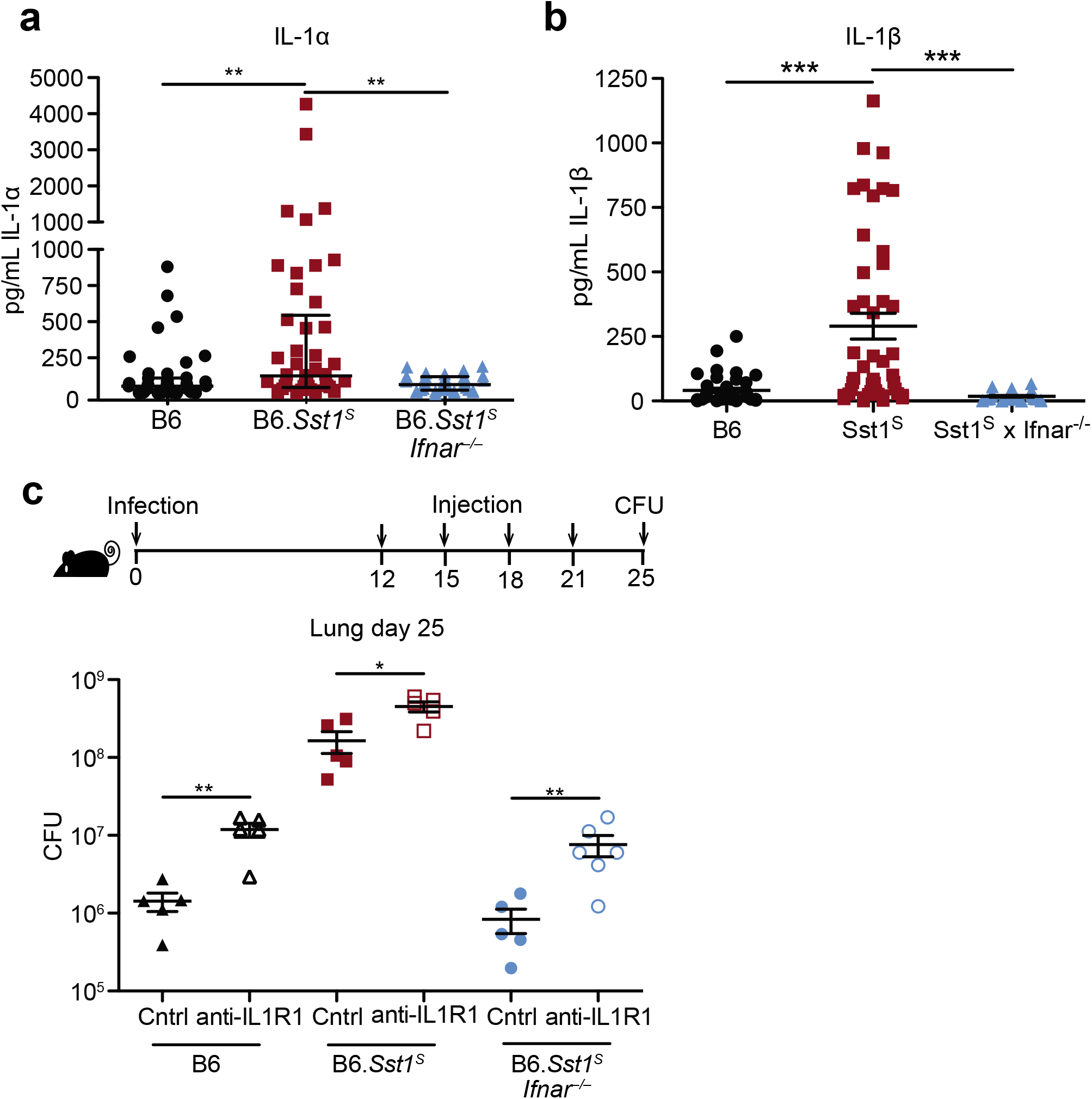
IFNAR signaling results in high but non-pathological IL-1 protein levels in B6.Sst1^S^ mice. **a-b**, Protein levels of IL-1α (**a**) and IL-1β (**b**) were measured in the lungs of Mtb-infected mice by ELISA at day 25. Combined results of four independent experiments. c, Lung bacterial burdens at day 25 from Mtb-infected mice treated with anti-IL1R1 or isotype control antibody. All animals except B6 were bred in-house (**a-c**). Error bars are SEM. Analyzed with two-ended Mann-Whitney test (**a-c**). Asterisk, *p* ≤ 0.05; two asterisks, *p* ≤ 0.01; three asterisks, *p* ≤ 0.001.

We reasoned that the high levels of IL-1α/β in B6.*Sst1^S^* mice may be a consequence of the higher bacterial burdens in these mice, or alternatively, may be causing increased bacterial replication via induction of a pro-bacterial inflammatory milieu, as previously proposed^38,39^. To distinguish these possibilities, we inhibited IL-1 signaling *in vivo* using an anti-IL-1R1 blocking antibody^40^, beginning 12 days post-infection (Fig. 2c, S4a). Both B6 and B6.*Sst1^S^* mice treated with IL-1R1 blocking antibody exhibited increased bacterial burdens compared to mice treated with an isotype control antibody. These results confirm prior evidence that IL-1 plays a protective role in B6 mice^41–46^, and extend this observation to B6.*Sst1^S^* mice as well. Thus, elevated IL-1 levels do not explain the exacerbated infections of B6.*Sst1^S^* mice.

Type I IFNs induce the expression of hundreds of target genes. However, given that IL-1 signaling is essential for resistance to *Mtb*^41,42^, we were particularly interested in *Il1rn*, which encodes the secreted IL-1 receptor antagonist (IL-1Ra)^47^. IL-1Ra binds to IL-1R1 without generating a signal and blocks binding of both IL-1α and IL-1β^48^ (Fig. 3a). *Il1rn* is known to be induced by type I IFN signaling^49,50^, and as expected, B6.*Sst1^S^* BMMs strongly upregulated *Il1rn* in an *Ifnar*-dependent manner when stimulated with TNF (Fig. 3b). Similarly, B6.*Sst1^S^* mice infected with *M. tuberculosis* had higher levels of IL-1Ra protein in their lungs, as compared to infected B6 or B6.*Sst1^S^Ifnar^−/−^* mice (Fig. 3c, S4b). These results raised the possibility that the high IL-1 protein levels in B6.*Sst1^S^* mice are inadequate to protect against infection because of a block in IL-1 signaling. To test this possibility, we examined the amount of functional IL-1 signaling in lung homogenates from infected mice using HEK-Blue-IL-1R^TM^ reporter cells (Invivogen). Despite higher levels of IL-1 proteins, lung homogenates from infected B6.*Sst1^S^* mice had less functional IL-1 signaling capacity as compared with B6 mice (Fig. 3d, S3c). The reporter appeared to be a reliable indicator of functional IL-1 as responses were blocked by anti-IL-1R1 antibody (Fig. S4c). The lower levels of IL-1 signaling seen in B6.*Sst1^S^* mice was reversed in B6.*Sst1^S^Ifnar^−/−^* mice. These data underline the importance of distinguishing IL-1 protein levels from signaling capacity, and suggest that B6.*Sst1^S^* mice may be susceptible to *Mtb* because of reduced functional IL-1 signaling, despite increased IL-1 protein levels. Consistent with these observations, some data suggest IL-1Ra may also exacerbate TB in humans^51–54^.

**Fig. 3.**
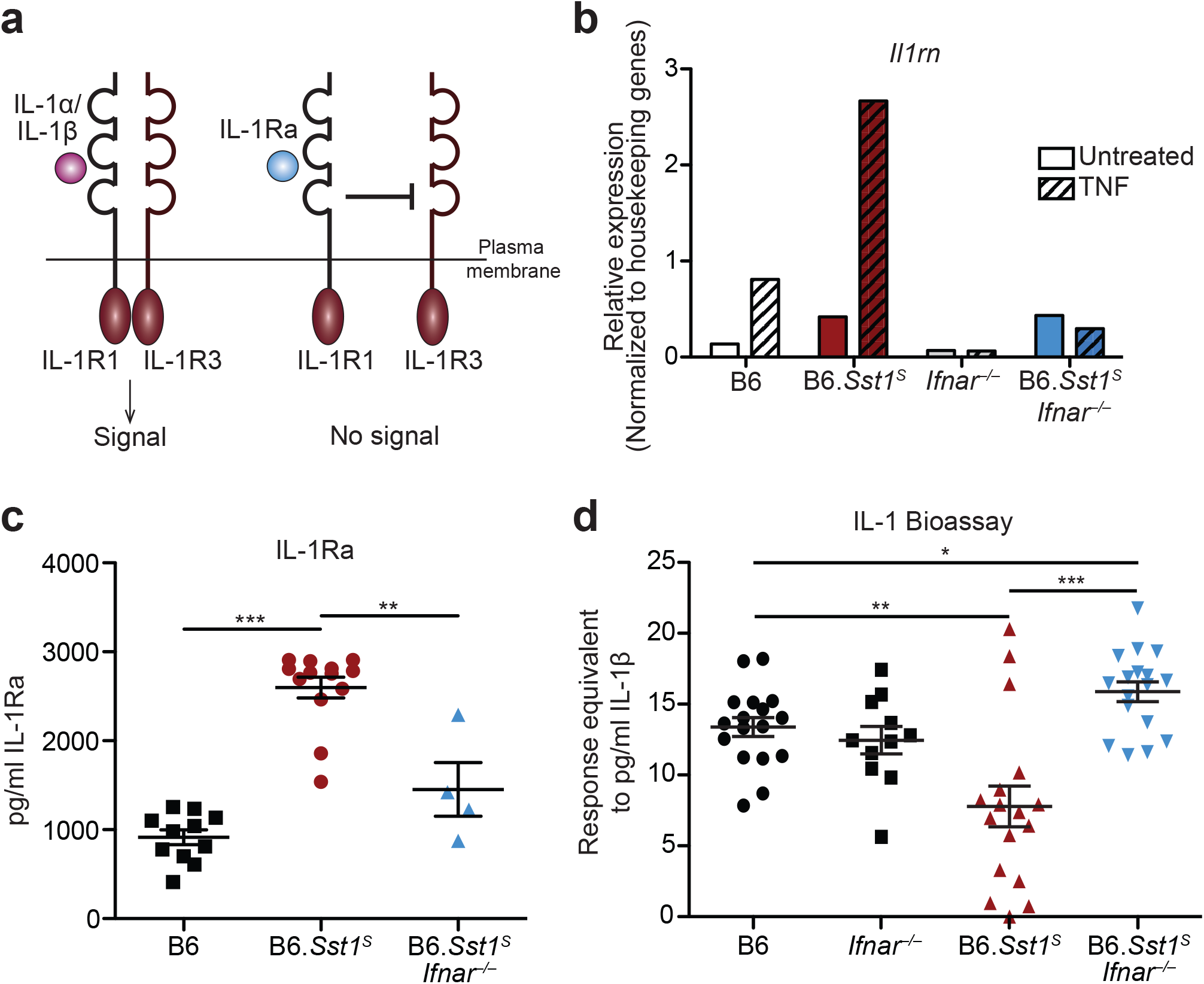
Elevated IL-1Ra and decreased functional IL-1 signaling in Mtb-infected B6.*Sst1^S^* mice. **a**, Schematic of IL-1R signaling. **b**, *Il1rn* expression in BMMs treated with 10 ng/ml TNFα for 24 hours. **c**, IL-1Ra protein levels and **d**, IL-1 reporter assay measuring functional IL-1 activity in lungs of Mtb-infected mice at day 25. Combined results of three independent infections (**d**) or representative of at least two independent experiments (**b, c**). All animals except B6 were bred in-house (**a-c**). Error bars are SEM. Analyzed with two-ended Mann-Whitney test (**c, d**). Asterisk, *p* ≤ 0.05; two asterisks, *p* ≤ 0.01; three asterisks, *p* ≤ 0.001.

To test whether excessive type I IFN signaling neutralizes IL-1 signaling via IL-1Ra, we sought to reduce IL-1Ra levels in B6.*Sst1^S^* mice during *Mtb* infection. To do this, we first crossed B6.*Sst1^S^* to *Il1rn^−/−^* mice^55^. Since uninfected homozygous *Il1rn* ^/^ mice exhibit signs of inflammatory disease due to dysregulated IL-1 signaling^55,56^, and because heterozygous *Il1rn^+/−^* mice have a partial decrease in IL-1Ra levels^55^, we generated both heterozygous B6.*Sst1^S^Il1rn^−/−^* and homozygous B6.*Sst1^S^Il1rn^−/−^* mice. Both heterozygous and homozygous *Il1rn* deficiency protected B6.*Sst1^S^* mice from *Mtb*. In fact, bacterial burdens in B6.*Sst1^S^Il1rn^−/−^* mice were even lower than those found in ‘resistant’ B6 mice (Fig. 4a). Notably, a partial reduction in IL-1Ra levels due to heterozygous deficiency of *Il1rn* was sufficient to almost entirely reverse the enhanced IFN-driven susceptibility of *Sst1^S^* mice (Fig. 4b). Histological samples of infected lungs showed significant reduction in lesion sizes in both B6.*Sst1^S^Il1rn^−/−^* and B6.*Sst1^S^Il1rn^−/−^* mice as compared to B6.*Sst1^S^* (Fig. 4c). Despite concerns that enhanced IL-1 signaling may cause immunopathology, both B6.*Sst1^S^Il1rn^−/−^* and B6.*Sst1^S^Il1rn^−/−^* mice retained more body weight and exhibited increased survival as compared to infected B6.*Sst1^S^* mice (Fig. 4b, Fig.S5a).

**Fig. 4.**
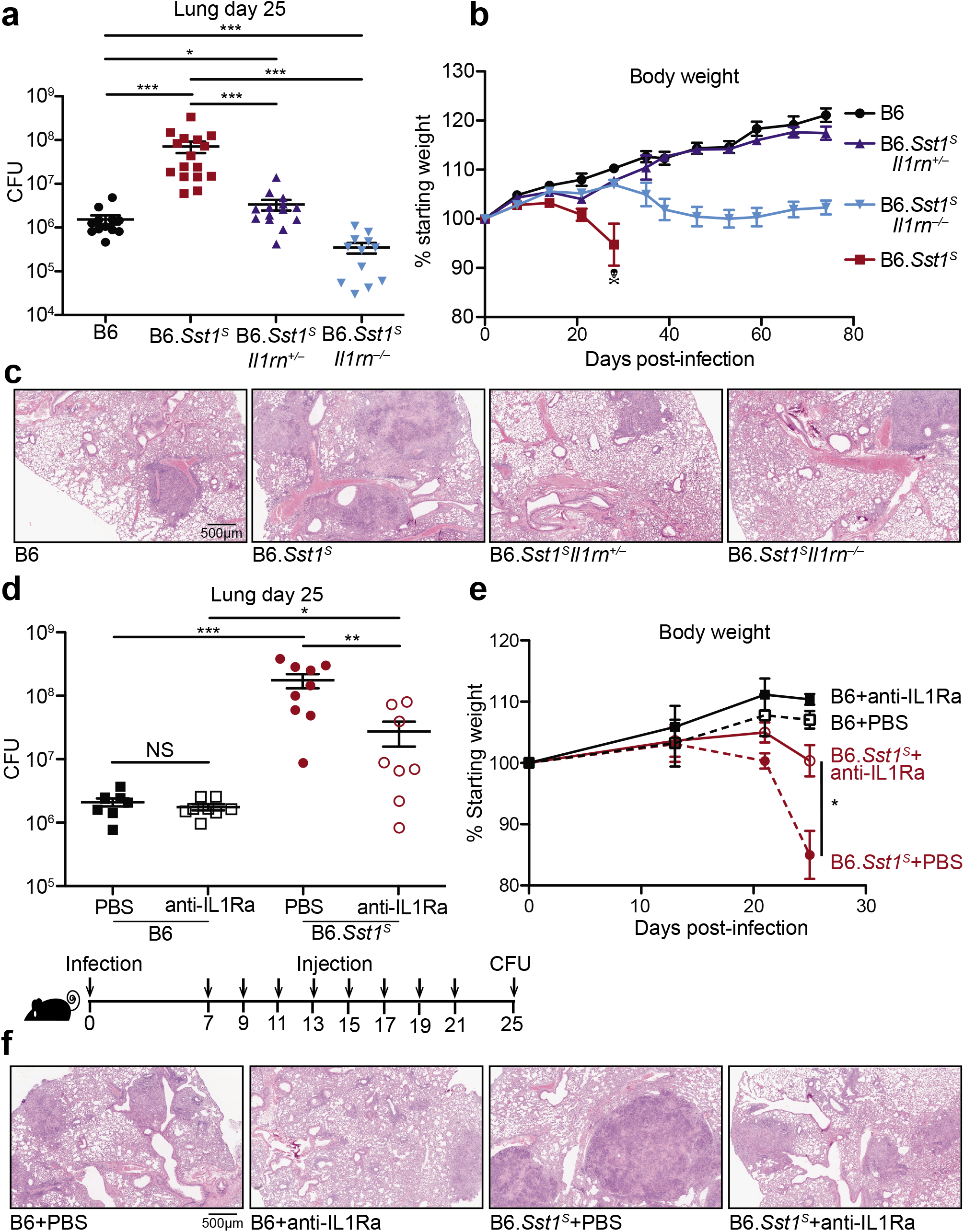
Inhibition of IL-1Ra rescues the susceptibility of B6.*Sst1^S^* mice. **a-c**, Lungs of Mtb-infected mice at day 25 were measured for (**a**) bacterial burdens in the lungs, (**b**) body weight, or (**c**) lung sections were stained with hematoxylin and eosin (H&E) for histology. d-f, Mtb-infected mice were treated with anti-IL1Ra antibody or PBS and (**d**) lung bacterial burdens were measured at day 25, or (**f**) day 25 lung sections were stained with H&E, or (**e**) body weights were recorded over time. All were bred in-house, and all except B6 and B6.*Sst1^S^* were littermates (**a-c**); B6.*Sst1^S^* were bred in-house (**d-f**). Error bars are SEM. Analyzed with two-ended Mann-Whitney test (**a, b, d, e**). Asterisk, *p* ≤ 0.05; two asterisks, *p* ≤ 0.01; three asterisks, *p* ≤ 0.001.

The dramatic protective effects of even partial reductions in IL-1Ra suggested that IL-1Ra might be a suitable target for host-directed therapy during *Mtb* infection. To test this idea, we treated infected B6.*Sst1^S^* mice with an anti-IL-1Ra antibody^57^ to block IL-1Ra and restore IL-1 signaling (Fig. 4d). B6.*Sst1^S^* mice that received the antibody for two weeks had significantly lower bacterial burdens in their lungs as compared to those receiving a control injection of PBS. In addition, mice treated with anti-IL-1Ra antibody retained significantly more body weight than their control counterparts (Fig. 4e), and exhibited reduced lung lesions (Fig. 4f), suggesting that the treatment did not cause detrimental inflammation. Overall these data indicate that genetic or antibody-mediated reduction of IL-1Ra rescues the type I IFN-driven susceptibility in B6.*Sst1^S^* mice without overt detrimental immunopathology.

## Discussion

There is increasing interest in developing host-directed therapeutics for Mtb^14^. Although such therapeutics have not yet proven to be curative, and may thus be unlikely to replace antibiotics, host-directed therapies may offer some advantages in specific scenarios. For example, host-directed therapy could serve as an adjunct to antibiotic regimens in multidrug-resistant or extensively-drug resistant tuberculosis, where mortality is at 20%^58,59^. It may also be more difficult for *Mtb* strains to evolve resistance to a host-directed therapy, an important consideration given the increasing prevalence of multi-drug resistant *Mtb* strains. In addition, our results, along with those of others^3,4^, suggest a strategy in which latently infected individuals could be screened for the type I IFN signature known to be a strong predictor of progression to active TB. If the type I IFN signature in these individuals could then be reset or reversed by a host-directed therapy targeting IL-1Ra, or other host factors^22^, then it might be possible to prevent progression to active TB — the transmissible form of the disease — without the prolonged (6-month) antibiotic treatments that are associated with poor compliance and emergence of drug-resistant strains. This therapeutic strategy, though only presented here as a proof-of-concept in animals, may help focus treatments on those who are likely to transmit among the billions of latently infected persons. The success of such a strategy would not necessarily require microbiological cure, but would depend on identification of highly potent master-regulators of interferon-driven TB disease that can be readily targeted clinically in TB endemic areas. Identification of such regulators and development of therapies appropriate for resource-limited settings will require better animal models of interferon-driven TB disease *in vivo*. Our results show that the B6.*Sst1^S^* mouse may represent a useful model of IFN-driven TB disease in humans and suggest that therapeutic targeting of IL-1Ra may be a potent strategy for reversing type I IFN-driven susceptibility *in vivo*.

## Acknowledgements

We thank the Stanley and Cox labs for discussions and for support with *Mtb* and BSL3 experiments. We thank the Barton lab for discussions, L. Flores, P. Dietzen and R. Chavez for technical assistance, H. Nolla and A. Valeros and the Cancer Research Laboratory for flow cytometry. We thank A. Sandstrom, I. Rauch and P. Mitchell for helpful discussions and technical support. We thank J. Price, L. DiPeso and R. Merl for support with the RNAseq. We thank B. Penn for comments on the manuscript. Generation of the anti-IL1RA antibody was supported by the Extramural Collaborative Research Program of Cancer Research Institute, Kanazawa University. K.H.D and R.E.V. were supported by Investigator in the Pathogenesis of Infectious Diseases awards from the Burroughs Wellcome Fund. R.E.V. is an HHMI Investigator and is supported by NIH grants AI075039 and AI066302.

## Methods

### Mice

All mice were specific pathogen-free, maintained under a 12-hr light-dark cycle (7AM to 7PM), and given a standard chow diet (Harlan irradiated laboratory animal diet) *ad libitum*. All mice were sex and age-matched at 6-8 weeks old at the beginning of infections. C57BL/6J (B6), B6.129S-*Il1rn^tm1Dih^*/J (*Il1rn^−/−^*), B6(Cg)-*Ifnar1^tm1.2Ees^*/J (*Ifnar^−/−^*) and B6.129S6-^*Ch25htm1Rus*^/J (*Ch25h^−/−^*) were originally purchased from Jackson Laboratories and subsequently bred at UC Berkeley. B6J.C3-*Sst1*^C3HeBFeJKrmn^ mice (referred to as B6.*Sst1^S^* throughout) were from the colony of I. Kramnik at Boston University and then transferred to UC Berkeley. *Sting^gt/gt^* mice were previously described^1^. All animals used in experiments were bred in-house unless otherwise noted in the figure legends. All animal experiments complied with the regulatory standards of, and were approved by, the University of California Berkeley Institutional Animal Care and Use Committee.

### Mycobacterium tuberculosis infections

*Mtb* strain Erdman (gift of S.A. Stanley) was used for all infections. Frozen stocks of this wild-type strain were made from a single culture and used for all experiments. Cultures for infection were grown in Middlebrook 7H9 liquid medium supplemented with 10% albumin-dextrose-saline, 0.4% glycerol and 0.05% Tween-80 for five days at 37°C. Mice were aerosol infected using an inhalation exposure system (Glas-Col, Terre Haute, IN). A total of 9ml of culture was loaded into the nebulizer calibrated to deliver ~20 to 50 bacteria per mouse as measured by colony forming units (CFUs) in the lungs 1 day following infection (data not shown). Mice were sacrificed at 14 days or 25 days post-infection to measure CFUs and/or cytokines. All but 1 lung lobe was homogenized in PBS plus 0.05% Tween-80 or processed for cytokines (see below), and serial dilutions were plated on 7H11 plates supplemented with 10% oleic acid, albumin, dextrose, catalase (OADC) and 0.5% glycerol. CFUs were counted 21 days after plating. The remaining lobe was used for histology or for RNA extraction. For histology the sample was fixed in 10% formalin for at least 48 hours then stored in 70% ethanol. Samples were sent to Histowiz Inc for embedding in wax, sectioning and staining.

### Cytokine measurements

Cell-free lung homogenates were generated as previously described^2^. Briefly, lungs were crushed through 100μm Falcon cell strainers in sterile PBS with 1% FBS and Pierce Protease Inhibitor (Thermo Fisher). An aliquot was removed for measuring CFU by plating as described above. Cells and debris were then removed by low-speed centrifugation (300×*g*) and the resulting cell-free homogenate was filtered twice with 0.2μm filters to remove all *Mtb* for work outside of BSL3. All homogenates were aliquoted, flash-frozen in liquid nitrogen and stored at –80°C. Each aliquot was thawed a maximum of twice to avoid potential artifacts due to repeated freeze-thaw cycles. All cytokines except IL-10 and IL-1Ra was measured using Cytometric Bead Assay (BD Biosciences) according to manufacturer protocols. Results were collected using BD LSRFortessa (BD Biosciences) and analyzed using FCAP Array v3.0. IL-10 levels were measured using Mouse IL-10 ELISA Ready-SET-Go! (2^nd^ Generation, eBioscience). IL-1Ra levels were measured by ELISA using Mouse IL-1ra/IL-1F3 Quantikine ELISA Kit (R&D Systems) according to manufacturer protocols.

### IL-1 bioactivity reporter assay

Mice were infected with ~25-35 bacteria per mouse and sacrificed at 25 days-post infection to prepare cell-free lung homogenates. Assays were performed using HEK-Blue™ IL-1R cells (InvivoGen) with minor modifications. Cells were cultured under antibiotic-selection according to manufacturer protocols. Cells were plated at 5×10^4^ cells/well in 50μl in 96-well plates, and allowed to adhere overnight in DMEM supplemented with 10% FBS, 2mM glutamine, 100 U/ml streptomycin and 100 μg/ml penicillin in a humidified incubator (37°C, 5% CO_2_). 50μl cell-free lung homogenates or standard curve made from recombinant mouse IL-1β (R&D systems 401-ML-005) were added to the cells and incubated overnight in a humidified incubator. Assays were developed using QUANTI-Blue (InvivoGen) according to manufacturer protocols. Experiments in which the initial CFU was either too high or too low produced inconsistent results in this assay.

### Flow cytometry

Lungs were perfused with 10 ml of cold PBS and strained through 40μm cell strainers. Aliquots were removed for quantifying CFU. Cells were washed and stained with fixable viability dye (Thermo Fisher 65-0865-14). An aliquot of cells from each sample were removed and mixed with counting beads (Thermo Fisher C36950) for later enumeration. The rest of the cells were incubated with anti-mouse CD16/CD32 monoclonal antibody to block Fc receptors (Thermo Fisher 14-0161-81), then with antibodies for surface staining. The following antigens were stained for: CD45 (30-F11, Biolegend 103107), CD11b (M1/70, Thermo Fisher 48-0112-82), CD11c (N418, Biolegend 117335), Ly6G (1A8, BD Biosciences 740554), Ly6C (HK1.4, Thermo Fisher 17-5932-80), CD24 (M1/69, BD Bioscience 564664), MHC II (M5/114.15.2, Biolegend 107625), SiglecF (E50-2440, BD Biosciences 562680). Cells were fixed with fixation buffer (BD Biosciences 554714) for at least 1 hour at room temperature and stored in PBS with 1% FBS and 2mM EDTA overnight at 4°C in the dark. Data were acquired on a BD Fortessa X-20 flow cytometer and analyzed with FlowJo v10.

### Bone marrow-derived macrophages (BMMs) and TNF-treatment

Bone marrow was harvested from mouse femurs and tibias, and cells were differentiated by culture on non-tissue culture-treated plates in RPMI supplemented with supernatant from 3T3-MCSF cells (gift of B. Beutler), 10% fetal bovine serum (FBS), 2mM glutamine, 100 U/ml streptomycin and 100 μg/ml penicillin in a humidified incubator (37°C, 5%CO_2_). BMMs were harvested six days after plating and frozen in 95% FBS and 5% DMSO. For *in vitro* experiments, BMMs were thawed into media as described above for 4 hours in a humidified 37°C incubator. Adherent cells were washed with PBS, counted and replated at 1.2 × 10^6^ ~ 1.5 × 10^6^ cells/well in a TC-treated 6-well plate. Cells were treated with 10 ng/ml recombinant mouse TNFα (410-TRNC-010, R&D systems) diluted in the media as described above.

### Quantitative RT-PCR

Total RNA from BMMs was extract using RNeasy total RNA kit (Qiagen) according to manufacturer specifications. Total RNA from infected tissues was extracted by homogenizing in TRIzol reagent (Life technologies) then mixing thoroughly with chloroform, both done under BSL3 conditions. Samples were then removed from the BSL3 facility and transferred to fresh tubes under BSL2 conditions. Aqueous phase was separated by centrifugation and RNA was further purified using an RNeasy total RNA kit (Qiagen). Equal amounts of RNA from each sample were treated with DNase (RQ1, Promega) and cDNA was made using Superscript III (Invitrogen). Complementary cDNA reactions were primed with poly(dT) for the measurement of mature transcripts. For experiments with multiple time points, samples were frozen in the RLT buffer (Qiagen) or RNAlater™ solution (Invitrogen). Quantitative PCR was performed using QuantiStudio 5 Real-Time PCR System (Applied Biosystems) with Power Sybr Green PCR Master Mix (Thermo Fisher Scientific) according manufacturer specifications.

Transcript levels were normalized to housekeeping genes *Rps17, Actb and Oaz1* unless otherwise specified. The following primers were used in this study. *Rps17* sense: CGCCATTATCCC CAGCAAG; *Rps17* antisense: TGTCGGGATCCACCTCAATG; *Oaz1* sense: GTG GTG GCC TCT ACA TCG AG; *Oaz1* antisense: AGC AGA TGA AAA CGT GGT CAG; *Actb* sense: CGC AGC CAC TGT CGA GTC; *Actb* antisense: CCT TCT GAC CCA TTC CCA CC; *Ifnb* sense: GTCCTCAACTGCTCTCCACT; *Ifnb* antisense: CCTGCAACCACCACTCATTC; *Il1rn* sense:

CGCCCTTCTGGGAAAAGACC, *Il1rn* antisense: CCGTGGATGCCCAAGAACAC; *Irf1* sense: TGAGGAAGGGAAGATAGCCG; *Irf1* antisense: TGTATGCCTATCCCAATGTCCC; *Irgm1* sense: AAAACCAGAGAGCCTCACCA; *Irgm1* antisense: ATGTTGGGGAGTAGTGGAGC; *Gbp4* sense: TGAGTACCTGGAGAATGCCCT; *Gbp4* antisense: TGGCCGAATTGGATGCTTGG; *Gbp5* sense: TGTTCTTACTGGCCCCTGCT; *Gbp5* antisense: CCAATGAGGCACAAGGGTTC; *Ifit3* sense: AGCCCACACCCAGCTTTT; *Ifit3* antisense: CAGAGATTCCCGGTTGACCT; *Stat1* sense:

CAGAAAAACGCTGGGAACAGA; *Stat1* antisense: CAAGCCTGGCTGGCAC; *Gbp7* sense: AGCAAGCCCAAGTTCACACT; *Gbp7* antisense: TCCGCTCTGTCAGTTCTGTG.

### RNA sequencing and analysis

Total RNA was isolated as described above. Illumina-compatible libraries were generated by the University of California, Berkeley, QB3 Vincent J. Coates Genomics Sequencing Laboratory. The libraries were multiplexed and sequenced using one flow cell on HiSeq4000 (Illumina) as 100bp paired-end reads. The data were aligned using Sleuth^3^ and analyzed using Kallisto^4^.

### Antibody-mediated neutralization

For all antibody treatments, the schedules are indicated in the figures. All treatments were delivered by intraperitoneal injection. Mouse anti-mouse IFNAR1 (MAR1-5A3) and isotype control (GIR208, mouse anti-human IFNGR-α chain) were purchased from Leinco Technologies Inc. Each mouse was given 500μg per injection. Hamster anti-IL1R1 antibody (mIL1R-M147) was obtained from Amgen. Isotype control was Ultra-LEAF Purified Armenian Hamster IgG Isotype Antibody from Biolegend (400940). Each mouse was given 200μg per injection. Armenian hamster anti-IL1Ra antibody was produced in-house using a previously published hybridoma line^5^. Cells were grown in Wheaton CELLine Bioreactor Flasks (Fisher Scientific) according to manufacturer instructions. Media in the cell compartment used ultra-low IgG FBS (ThermoFisher 16250078) to minimize bovine IgG contamination during purification. Cell-free supernatant from the cell compartment was purified using protein G resin (GenScript). IgG was eluted using 0.1M acetic acid, then 10% of total volume of 1M Tris pH8 and 0.5M NaCl was added to neutralize. Size exclusion and buffer exchange to PBS was performed using Amicon Ultra-4 Centrifugal Filter Units (EMD Millipore). The final product was filter sterilized and stored at –80°C. For injections antibody stocks were diluted in sterile PBS and each mouse received 500μg per injection.

### Statistical analysis

All survival data were analyzed with Log-rank (Mantel-Cox) Test. All other data were analyzed with Mann-Whitney test unless otherwise noted. Both tests were run using GraphPad Prism 5. Asterisk, *p* ≤ 0.05; two asterisks, *p* ≤ 0.01; three asterisks, *p* ≤ 0.001. All error bars are s.e.

---

**Fig. S1.**
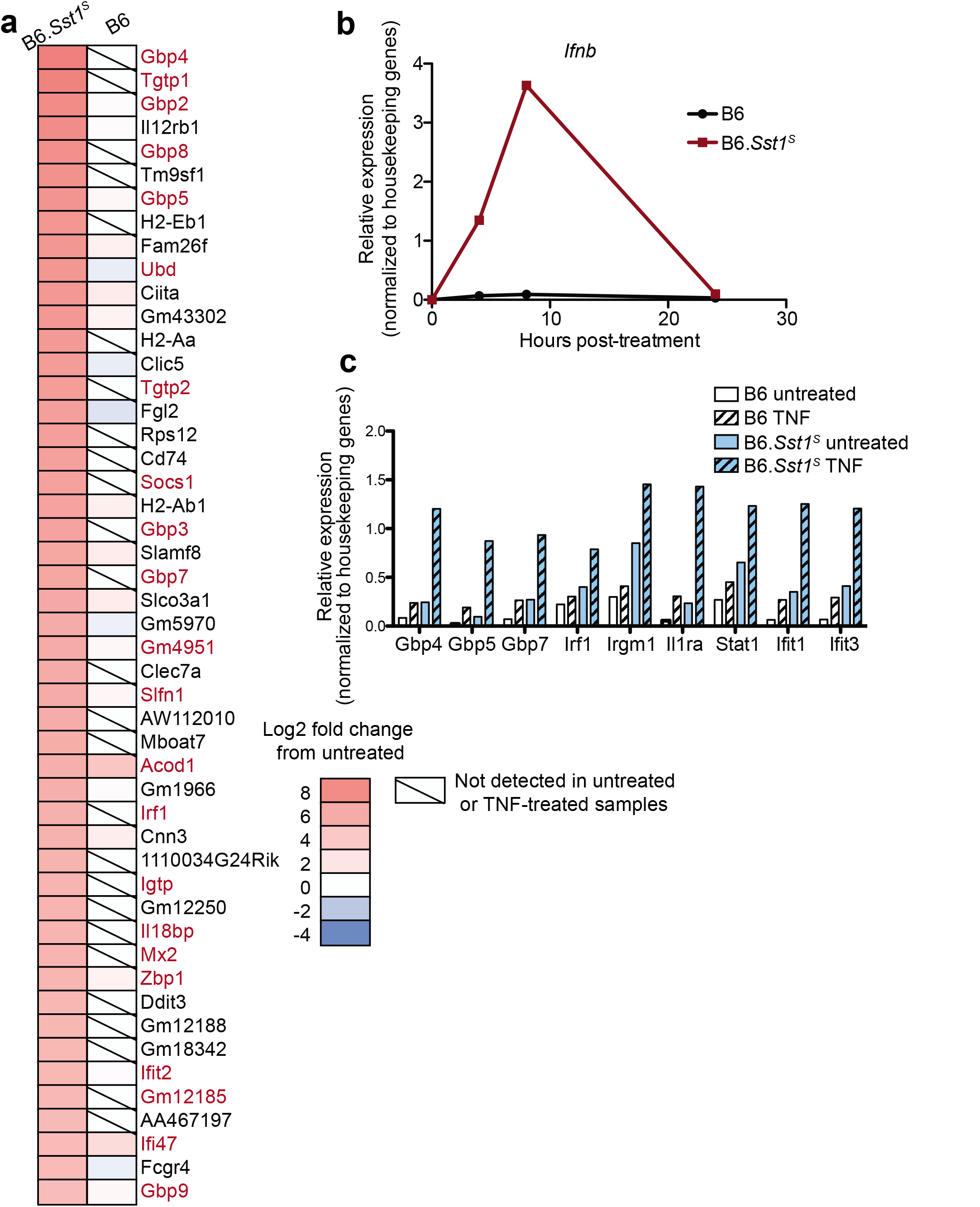
B6.*Sst1^S^* BMMs overexpress *Ifnb* and ISGs when stimulated with TNFα. **a**, RNAseq results showing fold upregulation of ISGs in TNFα -treated versus untreated BMMs. 50 most upregulated genes in B6.*Sst1^S^* BMMs are shown. Known ISGs names are marked in red. b-c, Expression of *Ifnb* (**b**) or selected ISGs (**c**) in BMMs measured by RT-qPCR. Results normalized to housekeeping genes. Representative data of at least two independent experiments.

**Fig. S2.**
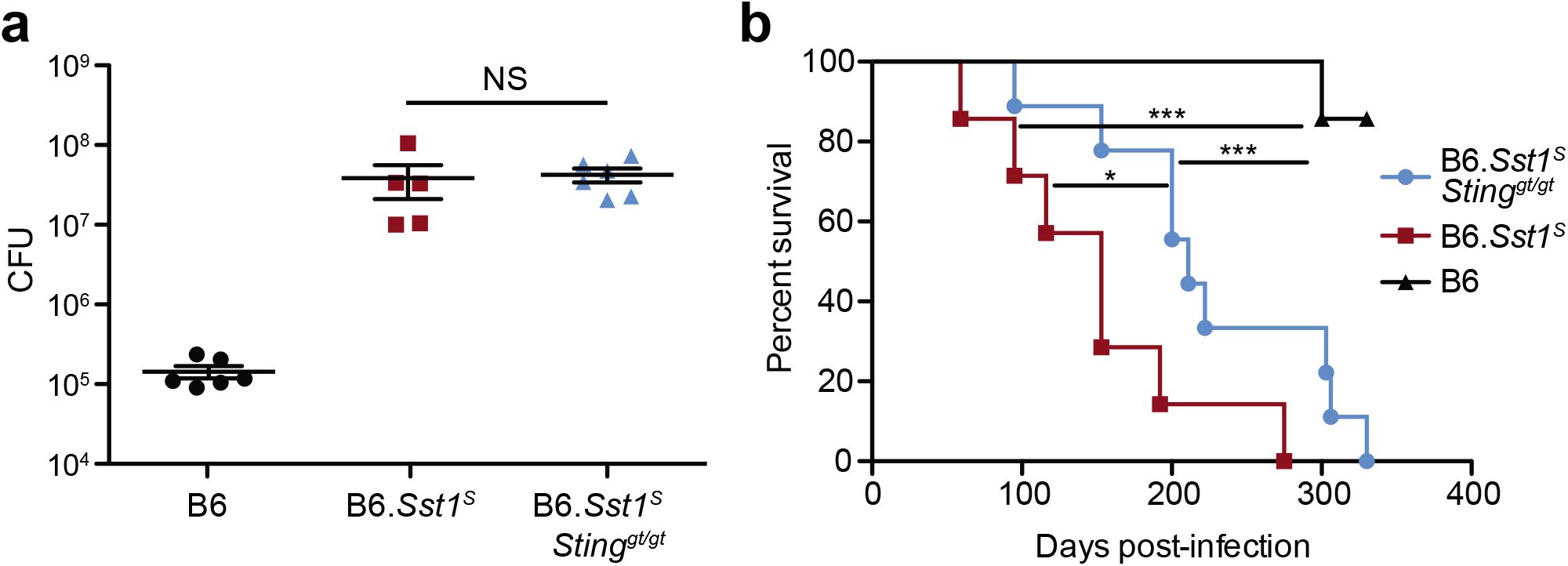
B6.*Sst1^S^Sting^gt/gt^* partially rescues the enhance susceptibility of B6.*Sst1^S^* mice to *Mtb*. Lung bacterial burdens at day 25 (representative of two independent infections) or **b**, Survival of mice infected with *Mtb*. All animals except B6 were bred in-house (**a-b**). Error bars are SEM. Analyzed with two-ended Mann-Whitney test (**a**) or Log-rank (Mantel-Cox) Test (**b**). Asterisk, *p* ≤ 0.05; two asterisks, *p* ≤ 0.01; three asterisks, *p* ≤ 0.001.

**Fig. S3.**
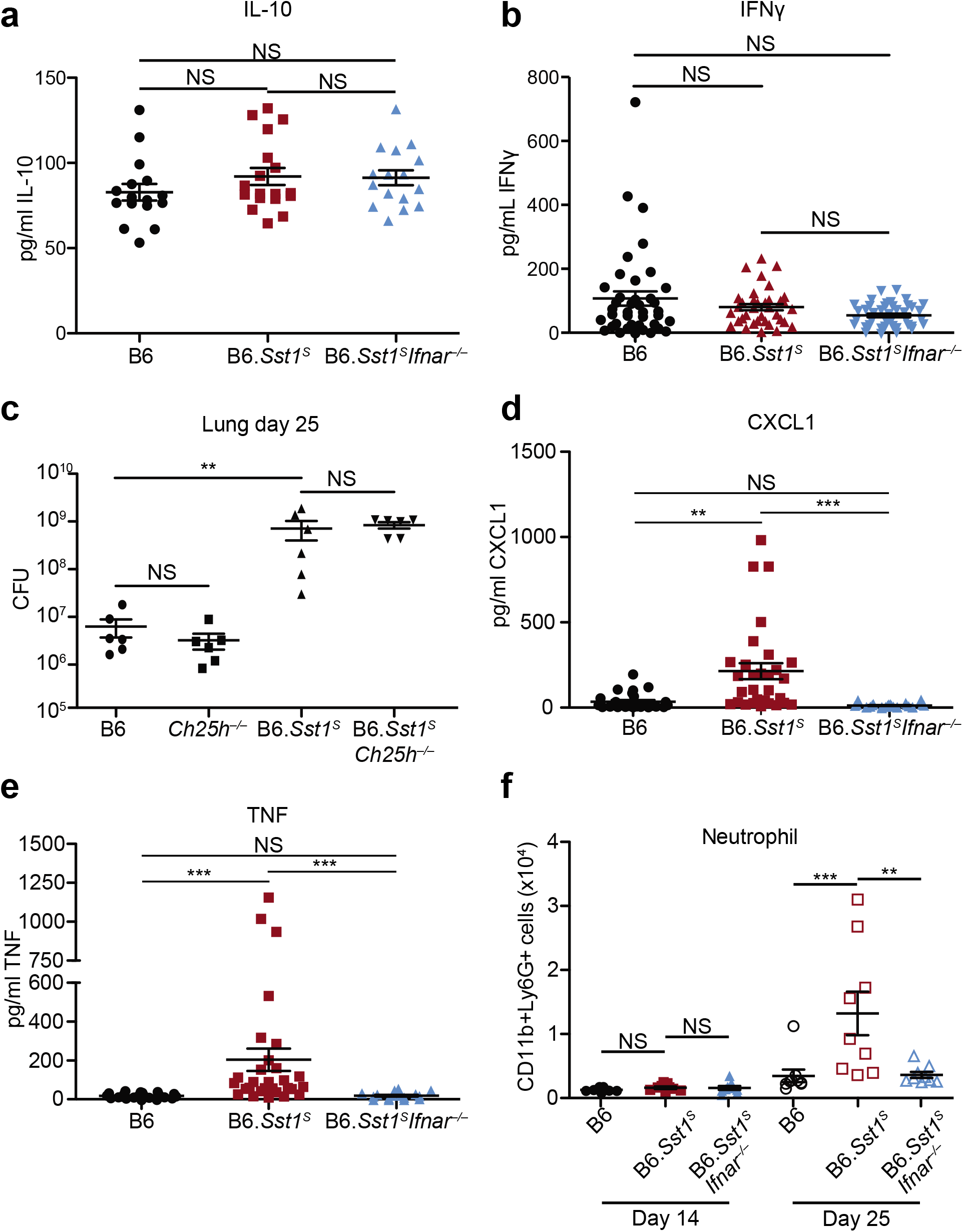
Enhanced inflammation in B6.*Sst1^S^* mice requires type I IFN but is not due to altered IL-10 or IFNγ levels or *Ch25h*. **a, b, d, e**, Protein levels of IL-10 (**a**), IFNγ (**b**), CXCL1 (**d**) and TNF (**e**) were measured in lungs of Mtb-infected mice at day 25. Combined results of three independent infections. **c**, Lung bacterial burden of Mtb-infected mice at day 25 (representative of two independent infections). **f**, Neutrophils (CD11b^+^Ly6G^+^) from lungs of Mtb-infected mice were enumerated on day 14 and day 25. Combined results of two independent infections. All animals except B6 were bred in-house (**a-e**); all animals were bred in-house (**f**). Error bars are SEM. Analyzed with two-ended Mann-Whitney test (**a-f**). Asterisk, *p* ≤ 0.05; two asterisks, *p* ≤ 0.01; three asterisks, *p* ≤ 0.001.

**Fig. S4.**
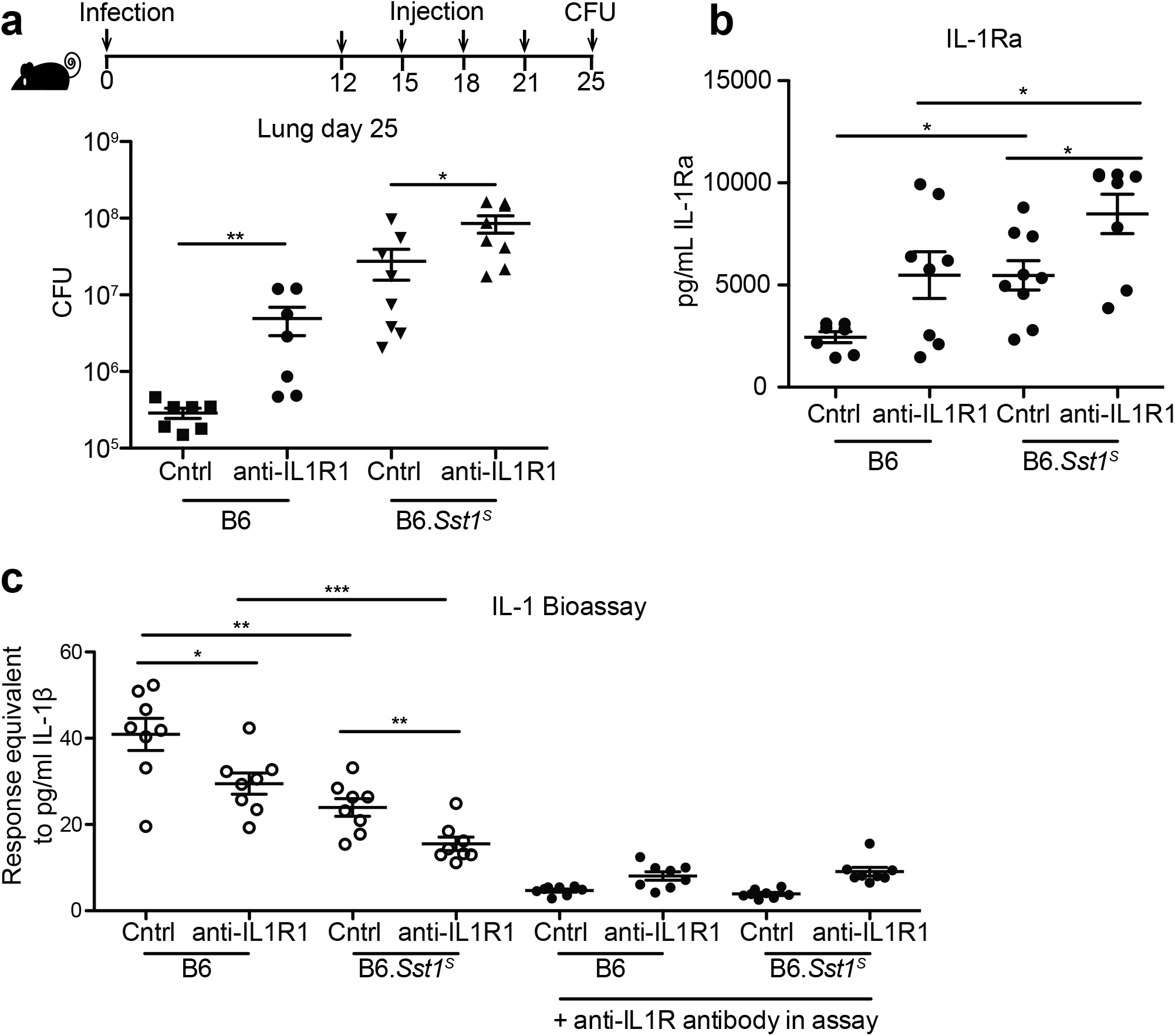
IL-1 blockade increases susceptibility in both B6 and B6.*Sst1^S^* mice. **a-c**, Mtb-infected mice were treated with anti-IL1R or isotype control antibodies, and on day 25 the lungs were measured for bacterial burden (**a**), IL-1Ra protein levels (**b**), and IL-1 bioactivity (**c**). All animals except B6 were bred in-house (**a-c**). Error bars are SEM. Analyzed with two-ended Mann-Whitney test (**a-c**). Asterisk, *p* ≤ 0.05; two asterisks, *p* ≤ 0.01; three asterisks, *p* ≤ 0.001.

**Fig. S5.**
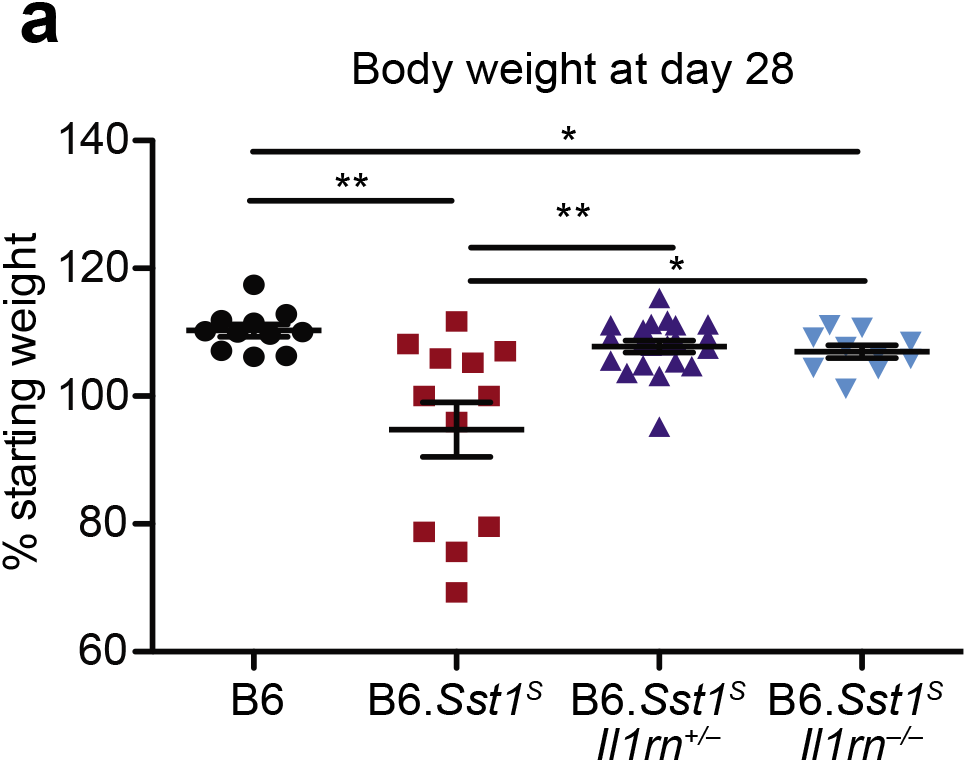
B6.*Sst1^S^* mice with homozygous or heterozygous *Il1rn* deletion retains more body weight at day 28 of *Mtb* infection than B6.*Sst1^S^* mice. Body weights on day 28 of individual mice shown in Fig. 4b. All were bred in-house, and all except B6 and B6. *Sst1^S^* were littermates. Error bars are SEM. Analyzed with two-ended Mann-Whitney test. Asterisk, *p* ≤ 0.05; two asterisks, *p* ≤ 0.01; three asterisks, *p* ≤ 0.001.

## References

1. World Health Organization. Global Tuberculosis Report 2017. Geneva; 2017.

2. Houben RM, Dodd PJ. The Global Burden of Latent Tuberculosis Infection: A Re-estimation Using Mathematical Modelling. PLoS Med. 2016;13(10):e1002152. doi:10.1371/journal.pmed.1002152.

3. Zak DE, Penn-Nicholson A, Scriba TJ, et al. A blood RNA signature for tuberculosis disease risk: a prospective cohort study. Lancet. 2016;387(10035):2312–2322. doi:10.1016/S0140-6736(15)01316-1.

4. Scriba TJ, Penn-Nicholson A, Shankar S, et al. Sequential inflammatory processes define human progression from M. tuberculosis infection to tuberculosis disease. PLoS Pathog. 2017;13(11):e1006687. doi:10.1371/journal.ppat.1006687.

5. Berry MP, Graham CM, Mcnab FW, et al. An interferon-inducible neutrophil-driven blood transcriptional signature in human tuberculosis. Nature. 2010;466(7309):973–977. doi:10.1542/peds.2011-2107LLLL.

6. Moreira-Teixeira L, Mayer-Barber K, Sher A, O’Garra A. Type I interferons in tuberculosis: Foe and occasionally friend. J Exp Med. 2018;215(5):1273–1285. doi:10.1084/jem.20180325.

7. Donovan ML, Schultz TE, Duke TJ, Blumenthal A. Type I Interferons in the Pathogenesis of Tuberculosis: Molecular Drivers and Immunological Consequences. Front Immunol. 2017;8(Nov):1633. doi:10.3389/fimmu.2017.01633.

8. Singhania A, Verma R, Graham CM, et al. A modular transcriptional signature identifies phenotypic heterogeneity of human tuberculosis infection. Nat Commun. 2018;9(2308). doi:10.1101/216879.

9. Pan H, Yan B-S, Rojas M, et al. Ipr1 gene mediates innate immunity to tuberculosis. Nature. 2005;434(7034):767–772. doi:10.1038/nature03419.

10. Pichugin AV, Yan BS, Sloutsky A, Kobzik L, Kramnik I. Dominant role of the sst1 locus in pathogenesis of necrotizing lung granulomas during chronic tuberculosis infection and reactivation in genetically resistant hosts. Am J Pathol. 2009;174(6):2190–2201. doi:10.2353/ajpath.2009.081075.

11. McNab F, Mayer-Barber K, Sher A, Wack A, O’Garra A. Type I interferons in infectious disease. Nat Rev Immunol. 2015;15(2):87–103. doi:10.1038/nri3787.

12. Boxx GM, Cheng G. The Roles of Type I Interferon in Bacterial Infection. Cell Host Microbe. 2016;19(6):760–769. doi:10.1016/j.chom.2016.05.016.

13. Nunes-Alves C, Booty MG, Carpenter SM, Jayaraman P, Rothchild AC, Behar SM. In search of a new paradigm for protective immunity to TB. Nat Rev Microbiol. 2014;12(4):289–299. doi:10.1038/nrmicro3230.

14. Hawn TR, Shah JA, Kalman D. New tricks for old dogs: Countering antibiotic resistance in tuberculosis with host-directed therapeutics. Immunol Rev. 2015;264(1):344–362. doi:10.1111/imr.12255.

15. Zhang G, deWeerd NA, Stifter SA, et al. A proline deletion in IFNAR1 impairs IFN-signaling and underlies increased resistance to tuberculosis in humans. Nat Commun. 2018;9(1):85. doi:10.1038/s41467-017-02611-z.

16. Manca C, Tsenova L, Freeman S, et al. Hypervirulent *M. tuberculosis* W/Beijing Strains Upregulate Type I IFNs and Increase Expression of Negative Regulators of the Jak-Stat Pathway. J Interf Cytokine Res. 2005;25(11):694–701. doi:10.1089/jir.2005.25.694.

17. Teles RMB, Graeber TG, Krutzik SR, et al. Type I Interferon Suppresses Type II Interferon-Triggered Human Anti-Mycobacterial Responses. Science (80-). 2013;339(6126):1448–1453. doi:10.1126/science.1233665.

18. Stanley SA, Johndrow JE, Manzanillo P, Cox JS. The Type I IFN response to infection with Mycobacterium tuberculosis requires ESX-1-mediated secretion and contributes to pathogenesis. J Immunol. 2007;178(5):3143–3152. doi:10.4049/jimmunol.178.5.3143.

19. Desvignes L, Wolf AJ, Ernst JD. Dynamic Roles of Type I and Type II IFNs in Early Infection with Mycobacterium tuberculosis. J Immunol. 2012;188(12):6205–6215. doi:10.4049/jimmunol.1200255.

20. Moreira-Teixeira L, Sousa J, McNab FW, et al. Type I IFN Inhibits Alternative Macrophage Activation during *Mycobacterium tuberculosis* Infection and Leads to Enhanced Protection in the Absence of IFN-γ Signaling. J Immunol. 2016;197(12):4714–4726. doi:10.4049/jimmunol.1600584.

21. Antonelli LR, Rothfuchs AG, Gonçalves R, et al. Intranasal poly-IC treatment exacerbates tuberculosis in mice through the pulmonary recruitment of a pathogen-permissive monocyte/macrophage population. J Clin Invest. 2010;120(5):1674–1682. doi:10.1172/JCI40817.

22. Mayer-Barber KD, Andrade BB, Oland SD, et al. Host-directed therapy of tuberculosis based on interleukin-1 and type I interferon crosstalk. Nature. 2014;511 VN-(7507):99–103. doi:10.1038/nature13489.

23. Dorhoi A, Yeremeev V, Nouailles G, et al. Type I IFN signaling triggers immunopathology in tuberculosis-susceptible mice by modulating lung phagocyte dynamics. Eur J Immunol. 2014;44(8):2380–2393. doi:10.1002/eji.201344219.

24. Bhattacharya B, Chatterjee S, Berland R, et al. Increased susceptibility to intracellular bacteria and necrotic inflammation driven by a dysregulated macrophage response to TNF. bioRxiv. 2018. http://biorxiv.org/content/early/2018/03/15/238873.abstract.

25. He X, Berland R, Mekasha S, et al. The sst1 Resistance Locus Regulates Evasion of Type I Interferon Signaling by Chlamydia pneumoniae as a Disease Tolerance Mechanism. PLoS Pathog. 2013;9(8). doi:10.1371/journal.ppat.1003569.

26. Dunn GP, Bruce AT, Sheehan KC, et al. A critical function for type I interferons in cancer immunoediting. Nat Immunol. 2005;6(7):722–729. doi:10.1038/ni1213.

27. Watson RO, Bell SL, MacDuff DA, et al. The Cytosolic Sensor cGAS Detects Mycobacterium tuberculosis DNA to Induce Type I Interferons and Activate Autophagy. Cell Host Microbe. 2014;17(6):811–819. doi:10.1016/j.chom.2015.05.004.

28. Wassermann R, Gulen MF, Sala C, et al. Mycobacterium tuberculosis Differentially Activates cGAS- and Inflammasome-Dependent Intracellular Immune Responses through ESX-1. Cell Host Microbe. 2015;17(6):799–810. doi:10.1016/j.chom.2015.05.003.

29. Wiens KE, Ernst JD. The Mechanism for Type I Interferon Induction by Mycobacterium tuberculosis is Bacterial Strain-Dependent. PLoS Pathog. 2016;12(8):1–20. doi:10.1371/journal.ppat.1005809.

30. Collins AC, Cai H, Li T, et al. Cyclic GMP-AMP Synthase Is an Innate Immune DNA Sensor for Mycobacterium tuberculosis. Cell Host Microbe. 2015;17(6):820–828. doi:10.1016/j.chom.2015.05.005.

31. Dey B, Dey RJ, Cheung LS, et al. A bacterial cyclic dinucleotide activates the cytosolic surveillance pathway and mediates innate resistance to tuberculosis. Nat Med. 2015;21(4):401–408. doi:10.1038/nm.3813.

32. McNab FW, Ewbank J, Howes A, et al. Type I IFN induces IL-10 production in an IL-27-independent manner and blocks responsiveness to IFN-γ for production of IL-12 and bacterial killing in Mycobacterium tuberculosis-infected macrophages. J Immunol. 2014;193(7):3600–3612. doi:10.4049/jimmunol.1401088.

33. Eshleman EM, Delgado C, Kearney SJ, Friedman RS, Lenz LL. Down regulation of macrophage IFNGR1 exacerbates systemic L. monocytogenes infection. PLoS Pathog. 2017;13(5):1–22. doi:10.1371/journal.ppat.1006388.

34. Reboldi A, Dang EV., McDonald JG, Liang G, Russell DW, Cyster JG. 25-Hydroxycholesterol suppresses interleukin-1 driven inflammation downstream of type 1 interferon. Science (80-). 2014;345(6197):679–684. doi:10.1126/science.1254790.

35. Novikov A, Cardone M, Thompson R, et al. Mycobacterium tuberculosis Triggers Host Type I IFN Signaling To Regulate IL-1 Production in Human Macrophages. J Immunol. 2011;187(5):2540–2547. doi:10.4049/jimmunol.1100926.

36. Mayer-Barber KD, Andrade BB, Barber DL, et al. Innate and Adaptive Interferons Suppress IL-1α and IL-1β Production by Distinct Pulmonary Myeloid Subsets during Mycobacterium tuberculosis Infection. Immunity. 2011;35(6):1023–1034. doi:10.1016/j.immuni.2011.12.002.

37. Mayer-Barber KD, Yan B. Clash of the Cytokine Titans: counter-regulation of interleukin-1 and type I interferon-mediated inflammatory responses. Cell Mol Immunol. 2016;14(April):1–14. doi:10.1038/cmi.2016.25.

38. Mishra BB, Rathinam VAK, Martens GW, et al. Nitric oxide controls the immunopathology of tuberculosis by inhibiting NLRP3 inflammasome-dependent processing of IL-1β. Nat Immunol. 2013;14(1):52–60. doi:10.1038/ni.2474.

39. Mishra BB, Lovewell RR, Olive AJ, et al. Nitric oxide prevents a pathogen-permissive granulocytic inflammation during tuberculosis. Nat Microbiol. 2017;2(May):17072. doi:10.1038/nmicrobiol.2017.72.

40. Nichols RD, Von Moltke J, Vance RE. NAIP/NLRC4 inflammasome activation in MRP8+ cells is sufficient to cause systemic inflammatory disease. Nat Commun. 2017;8(1). doi:10.1038/s41467-017-02266-w.

41. Fremond CM, Togbe D, Doz E, et al. IL-1 Receptor-Mediated Signal Is an Essential Component of MyD88-Dependent Innate Response to Mycobacterium tuberculosis Infection. J Immunol. 2007;179(2):1178–1189. doi:10.4049/jimmunol.179.2.1178.

42. Mayer-Barber KD, Barber DL, Shenderov K, et al. Caspase-1 Independent IL-1 Production Is Critical for Host Resistance to Mycobacterium tuberculosis and Does Not Require TLR Signaling In Vivo. J Immunol. 2010;184(7):3326–3330. doi:10.4049/jimmunol.0904189.

43. Yamada H, Mizumo S, Horai R, Iwakura Y, Sugawara I. Protective role of interleukin-1 in mycobacterial infection in IL-1 alpha/beta double-knockout mice. Lab Investig. 2000;80(5):759–767.

44. Sugawara I, Yamada H, Hua S, Mizuno S. Role of interleukin (IL)-1 type 1 receptor in mycobacterial infection. Microbiol Immunol. 2001;45(11):743–750.

45. Di Paolo NC, Shafiani S, Day T, et al. Interdependence between Interleukin-1 and Tumor Necrosis Factor Regulates TNF-Dependent Control of Mycobacterium tuberculosis Infection. Immunity. 2015;43(6):1125–1136. doi:10.1016/j.immuni.2015.11.016.

46. Juffermans NP, Florquin S, Camoglio L, et al. Interleukin-1 signaling is essential for host defense during murine pulmonary tuberculosis. J Infect Dis. 2000;182(3):902–908. doi:10.1086/315771.

47. Eisenberg S, Evans R, Arend W, et al. Primary structure and functional expression from complementary DNA of a human interleukin-1 receptor antagonist. Nature. 1990;343(6256):341–346. doi:10.1038/343341a0.

48. Dinarello CA. Overview of the IL-1 family in innate inflammation and acquired immunity. Immunol Rev. 2018;281(1):8–27. doi:10.1111/imr.12621.

49. Molnarfi N, Hyka-Nouspikel N, Gruaz L, Dayer J-M, Burger D. The production of IL-1 receptor antagonist in IFN-beta-stimulated human monocytes depends on the activation of phosphatidylinositol 3-kinase but not of STAT1. J Immunol. 2005;174(5):2974–2980. doi:10.4049/jimmunol.174.5.2974.

50. Corr M, Boyle DL, Ronacher LM, et al. Interleukin 1 receptor antagonist mediates the beneficial effects of systemic interferon beta in mice: implications for rheumatoid arthritis. Ann Rheum Dis. 2011;70(5):858–863. doi:10.1136/ard.2010.141077.

51. Janssen S, Schutz C, Ward A, et al. Mortality in Severe Human Immunodeficiency Virus-Tuberculosis Associates With Innate Immune Activation and Dysfunction of Monocytes. Clin Infect Dis. 2017;65(1):73–82. doi:10.1093/cid/cix254.

52. Settas LD, Tsimirikas G, Vosvotekas G, Triantafyllidou E, Nicolaides P. Reactivation of pulmonary tuberculosis in a patient with rheumatoid arthritis during treatment with IL-1 receptor antagonists (anakinra). J Clin Rheumatol. 2007;13(4):219–220. doi:10.1097/RHU.0b013e31812e00a1.

53. He D, Bai F, Zhang S, et al. High incidence of tuberculosis infection in rheumatic diseases and impact for chemoprophylactic prevention of tuberculosis activation during biologics therapy. Clin vaccine Immunol. 2013;20(6):842–847. doi:10.1128/CVI.00049-13.

54. Brassard P, Kezouh A, Suissa S. Antirheumatic drugs and the risk of tuberculosis. Clin Infect Dis. 2006;43(6):717–722. doi:http://dx.doi.org/10.1086/506935.

55. Hirsch E, Irikura VM, Paul SM, Hirsh D. Functions of interleukin 1 receptor antagonist in gene knockout and overproducing mice. Proc Natl Acad Sci U S A. 1996;93(20):11008–11013. doi:10.1073/pnas.93.20.11008.

56. Dinarello CA. Immunological and inflammatory functions of the interleukin-1 family. Annu Rev Immunol. 2009;27:519–550. doi:10.1146/annurev.immunol.021908.132612.

57. Fujioka N, Mukaida N, Harada A, et al. Preparation of specific antibodies against murine IL-1ra and the establishment of IL-1ra as an endogenous regulator of bacteria-induced fulminant hepatitis in mice. J Leukoc Biol. 1995;58(1):90–98.

58. Mitnick CD, Franke MF, Rich ML, et al. Aggressive regimens for multidrug-resistant tuberculosis decrease all-cause mortality. PLoS One. 2013;8(3):e58664. doi:10.1371/journal.pone.0058664.

59. Chung-Delgado K, Guillen-Bravo S, Revilla-Montag A, Bernabe-Ortiz A. Mortality among MDR-TB cases: comparison with drug-susceptible tuberculosis and associated factors. PLoS One. 2015;10(3):e0119332. doi:10.1371/journal.pone.0119332.

## Methods References

1. Sauer JD, Sotelo-Troha K, Von Moltke J, et al. The N-ethyl-N-nitrosourea-induced Goldenticket mouse mutant reveals an essential function of sting in the in vivo interferon response to Listeria monocytogenes and cyclic dinucleotides. Infect Immun. 2011;79(2):688–694. doi:10.1128/IAI.00999-10.

2. Mayer-Barber KD, Barber DL, Shenderov K, et al. Caspase-1 Independent IL-1 Production Is Critical for Host Resistance to Mycobacterium tuberculosis and Does Not Require TLR Signaling In Vivo. J Immunol. 2010;184(7):3326–3330. doi:10.4049/jimmunol.0904189.

3. Pimentel H, Bray NL, Puente S, Melsted P, Pachter L. Differential analysis of RNA-seq incorporating quantification uncertainty. Nat Methods. 2017;14(7):687–690. doi:10.1038/nmeth.4324.

4. Bray NL, Pimentel H, Melsted P, Pachter L. Near-optimal probabilistic RNA-seq quantification. Nat Biotechnol. 2016;34(5):525–527. doi:10.1038/nbt.3519.

5. Fujioka N, Mukaida N, Harada A, et al. Preparation of specific antibodies against murine IL-1ra and the establishment of IL-1ra as an endogenous regulator of bacteria-induced fulminant hepatitis in mice. J Leukoc Biol. 1995;58(1):90–98.

